# FISHFactor: A Probabilistic Factor Model for Spatial Transcriptomics Data with Subcellular Resolution

**DOI:** 10.1101/2021.11.04.467354

**Authors:** Florin C. Walter, Oliver Stegle, Britta Velten

## Abstract

**Motivation:** Factor analysis is a widely used tool for unsupervised dimensionality reduction of high-throughput data sets in molecular biology, with recently proposed extensions designed specifically for spatial transcriptomics data. However, these methods expect (count) matrices as data input and are therefore not directly applicable to single molecule resolution data, which are in the form of coordinate lists annotated with genes and provide insight into subcellular spatial expression patterns. To address this, we here propose FISHFactor, a probabilistic factor model that combines the benefits of spatial, non-negative factor analysis with a Poisson point process likelihood to explicitly model and account for the nature of single molecule resolution data. In addition, FISHFactor shares information across a potentially large number of cells in a common weight matrix, allowing consistent interpretation of factors across cells and yielding improved latent variable estimates.

**Results:** We compare FISHFactor to existing methods that rely on aggregating information through spatial binning and cannot combine information from multiple cells, and show that our method leads to more accurate results on simulated data. We demonstrate on a real data set that FISHFactor is able to identify major subcellular expression patterns and spatial gene clusters in a data-driven manner.

**Availability and Implementation:** The model implementation, data simulation and experiment scripts are available under https://www.github.com/bioFAM/FISHFactor.

## 1 Introduction

Transcriptomic profiling of individual cells using single-cell RNA sequencing is now a widely accessible tool for studying cellular heterogeneity in tissues and has contributed to the discovery of new cell types. However, single-cell RNA sequencing protocols are based on a disassociation step and therefore can provide only limited insight into the spatial organization of tissue and no information at all about the localization of RNA molecules within a cell. To address this, a growing number of spatially resolved transcriptomic technologies are being developed that allow measurements of gene expression while retaining spatial context (Rao *et al*., 2021; Palla *et al*., 2022). For example, next-generation sequencing coupled with spatial barcodes provides whole transcriptome measurements of tissue regions (Ståhl *et al*., 2016; Rodriques *et al*., 2019), but the resolution of current methods is at most at the level of individual cells and cannot resolve subcellular patterns. On the other hand, imaging-based techniques such as in-situ sequencing (Ke *et al*., 2013; Lee *et al*., 2015; Wang *et al*., 2018; Chen *et al*., 2018) or fluorescence in-situ hybridization (FISH) achieve subcellular resolution by measuring spatial positions of individual molecules. While FISH technologies were originally limited to the detection of a single or at most a handful of genes (Femino *et al*., 1998; Raj *et al*., 2008; Lyubimova *et al*., 2013), advances in imaging technologies, sequential hybridization and barcoding strategies nowadays enable probing tens to thousands of genes in a single experiment (Codeluppi *et al*., 2018; Lubeck and Cai, 2012; Lubeck *et al*., 2014; Chen *et al*., 2015; Eng *et al*., 2017, 2019), thus rendering such techniques increasingly applicable for the identification of subcellular gene expression patterns at scale.

Despite the availability of technologies that provide single-molecule resolution, most established analysis strategies for processing these data do not fully exploit the given resolution. Instead, RNA quantifications are limited to cellular resolution, for example by aggregating the numbers of molecules per cell (Eng *et al*., 2019; Chen *et al*., 2015; Codeluppi *et al*., 2018), or used only for the task of cell type inference or clustering (Park *et al*., 2021; Littman *et al*., 2021; Qian *et al*., 2020; Partel and Wählby, 2021). Thereby, such approaches cannot model subcellular gene expression patterns, which can provide important insights into cellular states, heterogeneity within cell types (Xia *et al*., 2019; Buxbaum *et al*., 2015) and can modulate the function of genes (Eng *et al*., 2019). Bento (Mah *et al*., 2022) was recently proposed as a tool to predict subcellular gene expression patterns and extract localization signatures. However, it requires the allowed spatial patterns to be predefined. With the increasing throughput of single-molecule techniques, it will become ever more important to identify major subcellular gene expression patterns in a data driven manner and use them as additional source of information when dissecting cell-to-cell heterogeneity.

Factor models are already widely used for the unsupervised discovery of the principal sources of variation in high-dimensional molecular data sets (Stein-O’Brien *et al*., 2018; Brunet *et al*., 2004; Witten *et al*., 2009; Argelaguet *et al*., 2018, 2020; Risso *et al*., 2018), and recent extensions to spatial data have successfully identified spatial gene expression patterns at the cellular level (Velten *et al*., 2022; Townes and Engelhardt, 2021; Berglund *et al*., 2018). However, these methods cannot leverage the subcellular resolution of spatial transcriptomics data, as they require a count matrix as input and consequently are not directly applicable to single molecule resolved data, which are lists of coordinates annotated with gene labels. To apply these methods, it is currently required to crudely aggregate the data, using spatial binning or summation of molecules per cell, which involves additional parameters and results in a loss of the exact spatial information.

To address these shortcomings, we here propose FISHFactor, a principled factor analysis framework that opens up the application of factor models for spatially resolved single-molecule data and enables the unbiased identification and discovery of subcellular expression patterns (Fig. 1). Other than existing spatial factor models, FISHFactor employs spatial Poisson point processes as observation model to explicitly model the subcellular coordinates of each RNA molecule. It can thereby fully leverage the single-molecule resolution of the data. We combine this with a spatially aware inference of factors using Gaussian processes tailored to spatial transcriptomics data and impose interpretable factors and weights using non-negativity constraints. To enable the integration and comparison of subcellular localization patterns across a population of cells, FISHFactor jointly models the information from multiple cells in a scalable manner by inferring a shared weight matrix, while retaining independent sets of factors. We assess the model using simulated data, where we demonstrate advantages of FISHFactor over existing approaches that require spatial binning and show the benefit of jointly modeling multiple cells. Using a real data set, we illustrate the use of FISHFactor to reveal subcellular localization patterns of genes and to analyze the co-localization of genes within a cell.

**Figure 1:**
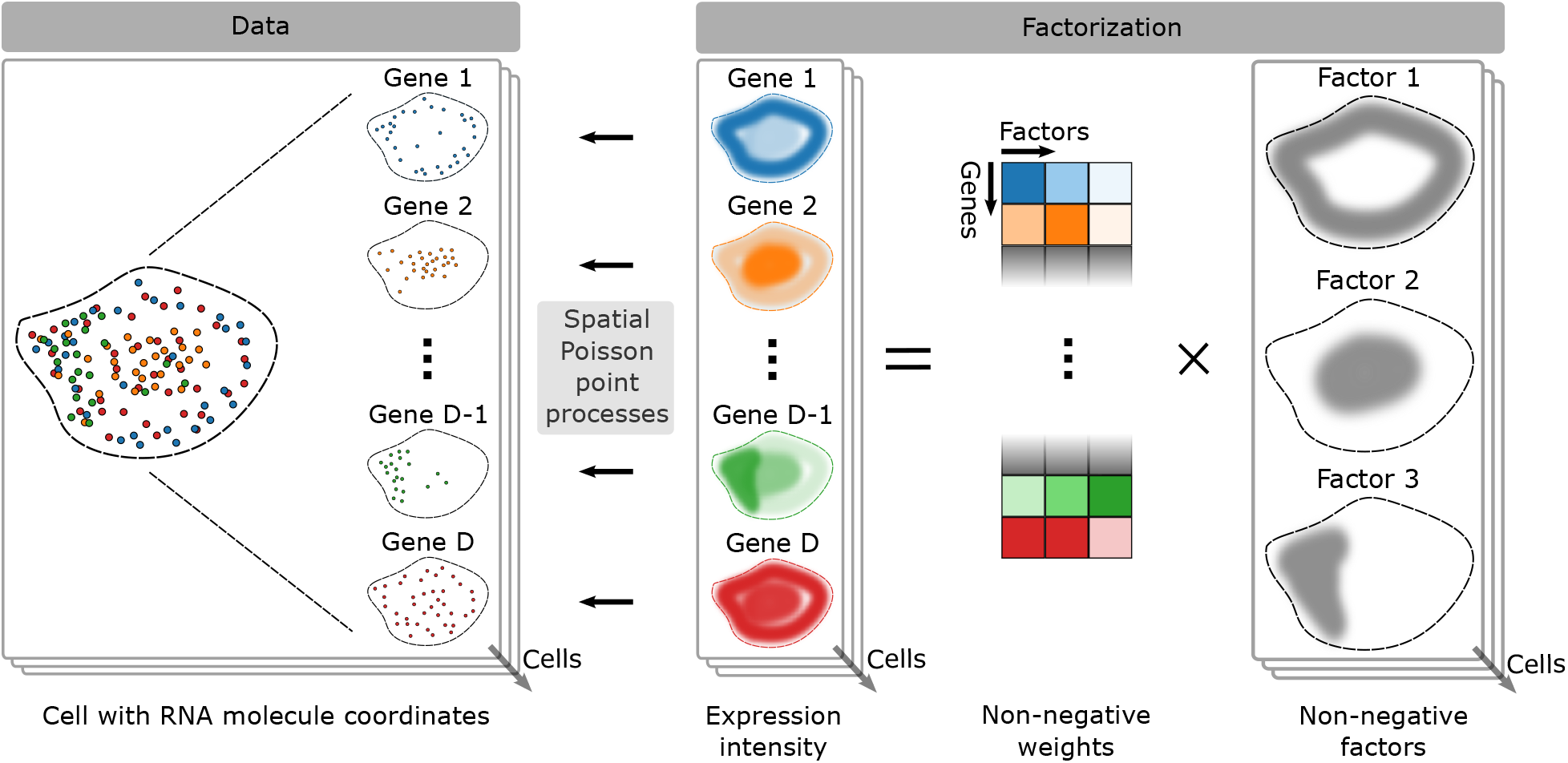
Illustration of FISHFactor for the analysis of spatial transcriptomics data with subcellular resolution. The input dataset (left) consists of RNA molecule coordinates in a single or optionally multiple segmented cells, e. g. from multiplexed FISH measurements. FISHFactor models the observed coordinates as realizations of a spatial Poisson point process with a gene-wise intensity function in each cell. The intensity functions of all genes and cells are governed by a low-dimensional decomposition into non-negative spatially-aware factors and non-negative weights (right, illustrated for 3 latent factors). Weights are shared between cells, whereas factors are specific to individual cells.

## 2 Materials and methods

### 2.1 Factor analysis for count data

Factor analysis is a dimensionality reduction technique commonly used for unsupervised analysis of highdimensional omics data sets (Stein-O’Brien *et al*., 2018). Based on correlation structures in a high-dimensional feature space, the method aims to find a low-dimensional embedding in terms of a small number of latent factors, representing the major axes of variation in the data. Starting from a high-dimensional dataset 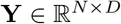 with *N* observations of *D* features, factor analysis finds a factorization **Y** = **ZW***^T^* + **E** with *K* latent factors 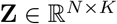 (typically *K* ≪ D), associated factor weights 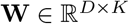 and residual noise 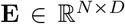. In contrast to non-linear dimensionality reduction methods, factor analysis identifies latent embeddings that can be directly interpreted, because the weights linearly link each latent factor to molecular features. Formulated in a probabilistic framework, factor analysis further allows for the incorporation of prior knowledge and various sparsity assumptions through the use of appropriate prior distributions, provides uncertainty estimates for the inferred variables and can account for different data types through the use of appropriate likelihood models. As a baseline model for sequencing data we here consider a Poisson likelihood, which can account for the count nature of the data and has been successfully applied to transcriptomics data (Townes *et al*., 2019). The decomposition in a Poisson factor model is given by

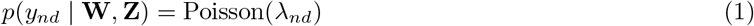

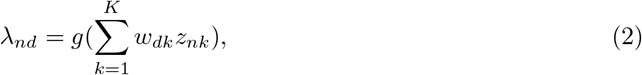

where *g* denotes a positive inverse-link function such as the exponential or softplus.

### 2.2 Non-negative factor analysis

To improve the interpretability and identifiability of factor analysis, different sparsity assumptions on the factors and weights have been employed, including sparsity on the level of features or sets of features (Witten *et al*., 2009; Argelaguet *et al*., 2018, 2020) as well as non-negativity constraints (Lee and Seung, 1999; Townes and Engelhardt, 2021). The latter have been found particularly useful, as they allow to find additive nonnegative spatial patterns and molecular signatures. In practice, non-negativity is achieved by constraining weights and factors to non-negative values, for example using non-negative parametrization or non-negative priors. In such a model, with Gaussian priors on the unconstrained latent variables, *λ_nd_* in Eq. 1 is given by

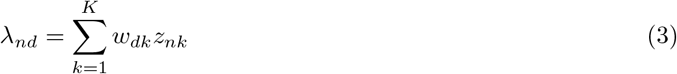

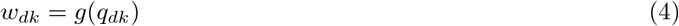

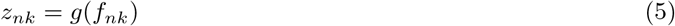

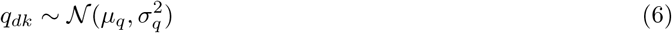

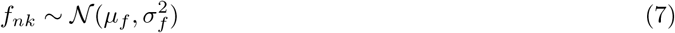

where *g* is a positive link-function and the parameters in Eq. 6, Eq. 7 are fixed hyperparameters.

### 2.3 Gaussian process factor analysis

A limitation of classical factor models in applications to spatial data is the assumption of independent observations *n* = 1,…, *N*. While this assumption may be appropriate for some data types, it generally does not hold for spatial data, where each observation comes with a spatial coordinate and spatial structures are present between samples. For example, gene expression profiles at nearby points are expected to be more similar than at points that are far apart. This spatial covariance can be incorporated into factor analysis by replacing the univariate Gaussian priors on factors in Eq. 7 by multivariate priors that can model covariation across samples. A flexible choice for this purpose are Gaussian process (GP) priors, which provide a nonparametric framework to model continuous dependencies between samples. This has given rise to Gaussian process factor analysis (GPFA) (Yu *et al*., 2008), where independent GP priors are placed on the factors to model smooth temporal patterns. The same concept has recently been applied for the identification of patterns in spatial transcriptomics data, in combination with different likelihood models and sparsity constraints (Velten *et al*., 2022; Townes and Engelhardt, 2021). In particular, this approach corresponds to replacing the factor prior in Eq. 7 with a Gaussian process prior:

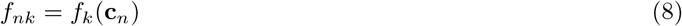

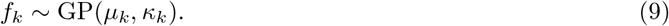

Here, 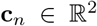 is the spatial coordinate of sample *n, μ_k_* is a mean function in 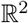, and *κ_k_* is a kernel function in 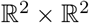. The choice of the kernel function determines the covariance structure. For example, a squared exponential kernel generates very smooth patterns, whereas a Matérn kernel leads to a more angular appearance as often observed for spatial expression patterns (Townes and Engelhardt, 2021).

### 2.4 Poisson point process likelihood

In contrast to (spatial) transcriptomics data sets at the cellular level, single-molecule resolved data consists of a list of *N* coordinate vectors {**c**_*n*_}_*n*=1…,*N*_ with 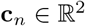 with gene annotations. For such data, the Poisson likelihood used in the discussed models can only be employed after a pre-processing step that aggregates the number of molecules in a certain spatial region or cell and ignores the exact spatial information. A more suitable likelihood model for single-molecule resolved data are Poisson point processes, which directly model the coordinates of each molecule. Poisson point processes have already been successfully used in GPFA with temporal data in neuroscience (Duncker and Sahani, 2018) and for cell typing in spatial data (Qian *et al*., 2020) but so far have not been considered in factor models for spatial transcriptomics data. Formally, an inhomogeneous spatial Poisson point process is characterized by a non-negative intensity function 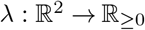. A set of point coordinates then has the probability density

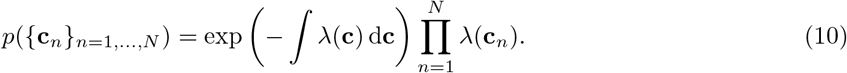

Intuitively, this means that more points are expected in regions where λ is high, and vice versa.

### 2.5 The FISHFactor Model

FISHFactor is a probabilistic factor model for single-molecule resolved spatial transcriptomics data that combines the concepts discussed in the preceding sections (Fig. 1): (i) Spatially aware inference of factors using Gaussian processes, (ii) interpretable factors and weights using non-negativity constraints and (iii) a likelihood model accounting for the nature of single-molecule data using inhomogeneous Poisson point processes. In addition, FISHFactor allows to integrate and compare inferred patterns across multiple cells by inferring a shared weight matrix.

The input data to FISHFactor consists of a list of spatial molecule coordinates 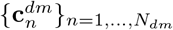 with 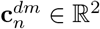 for a set of genes *d* = 1,…, *D* and cells *m* = 1,…, *M*. The assignment of molecules to cells is assumed to be known and can be defined from the image using existing segmentation techniques (Littman *et al*., 2021; Petukhov *et al*., 2022). FISHFactor models the coordinates as realizations of spatial Poisson point processes, where the gene- and cell-wise intensity functions *λ_dm_* are given by a decomposition into cell-specific factors and a weight matrix that is shared between cells. The generative model of FISHFactor is defined as

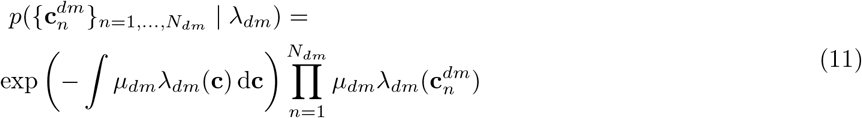

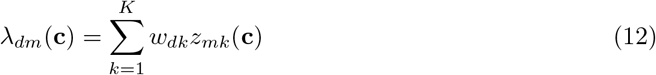

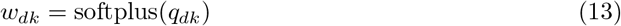

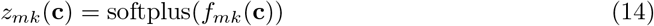

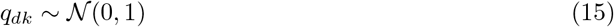

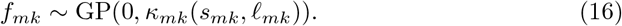

The integration limits in Eq. 11 are given by the respective cell boundaries, which are estimated by thresholding a kernel density estimate based on all associated molecules. The average intensity per gene and cell *μ_dm_* serves as a scale factor for the intensity function *λ_dm_* to account for differences in the overall expression intensities of genes and cells, to ensure that the inferred latent variables do not reflect abundances but sub-cellular patterns. It is determined as the number of molecules *N_dm_* divided by the cell area. The GP prior in Eq. 16 uses a Matérn kernel *κ_mk_* with smoothness parameter *v* and learnable output and length scales *s_mk_* and *ℓ_mk_*. The graphical model of FISHFactor is shown in Fig. 2.

**Figure 2:**
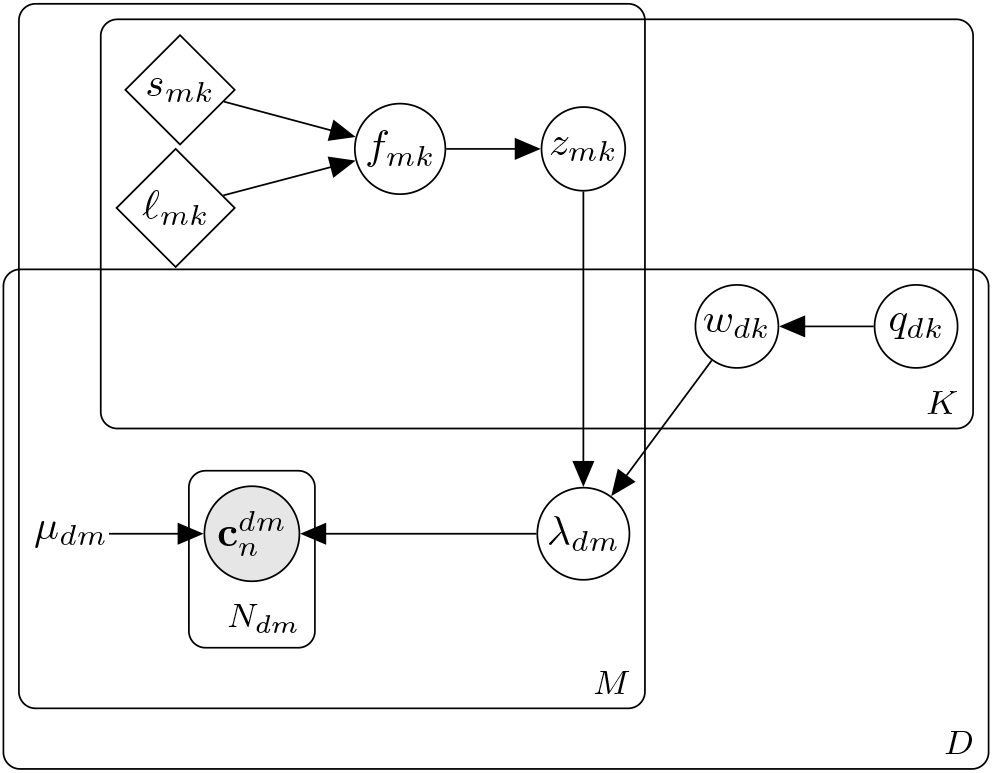
Graphical model of FISHFactor with *K* factors, *D* genes, *M* cells and *N_dm_* molecules per gene and cell. Gray nodes indicate observed variables, white nodes latent variables, rhombuses learnable parameters, and *μ_dm_* is a constant determined by the data.

### 2.6 Implementation

To infer the model’s latent variables in a scalable manner, FISHFactor is implemented using stochastic variational inference (Hoffman *et al*., 2013) and sparse approximations of the Gaussian processes (Hensman *et al*., 2015). In addition, a sequential update of cell-wise parameters is used to keep memory requirements constant in the number of cells, which otherwise can be a major bottleneck to the application of models to many cells. For this, every epoch consists of as many optimization steps as there are cells, whereby in every step one cell is loaded into memory, its parameters are optimized and the global weight matrix is updated.

Optimization of the evidence lower bound (ELBO) is performed with a clipped adam optimizer and a learning rate of 5 · 10^-3^. To determine convergence, the ELBO is monitored for each cell and the optimization is terminated as soon as it does not increase by a given value for any of the cells in a given number of epochs. FISHFactor is implemented using the probabilistic programming language Pyro (Bingham *et al*., 2019) and the low-level Pyro interface of GPyTorch (Gardner *et al*., 2018).

## 3 Results

### 3.1 FISHFactor outperforms existing factor models on simulated data

First, we validated FISHFactor’s ability to infer subcellular expression patterns on simulated data for individual cells (*M* = 1) and compared its performance to related existing factor model implementations. We considered non-negative matrix factorization (NMF) (Pedregosa *et al*., 2011), a widely-used method for a non-negative decomposition without spatial awareness, and non-negative spatial factorization (NSF) (Townes and Engelhardt, 2021), a recently proposed Gaussian process factor model for a non-negative decomposition with spatial awareness and a Poisson observation model. In contrast to FISHFactor, both methods require aggregation of molecule coordinates in spatial bins to obtain count matrices, for which we included different binning resolutions in the comparison (2 × 2, 5 × 5, 10 × 10, 20 × 20, 30 × 30, 40 × 40 and 50 × 50).

Data was simulated in form of molecule coordinates for 20 cells, where for each cell we independently simulated subcellular expression patterns following Eq. 12 for 50 genes using 3 latent spatial factors and corresponding gene weights and then sampled molecule coordinates from the resulting intensity function according to a spatial Poisson point processes by thinning (Lewis and Shedler, 1979). Cell shapes were obtained from real data (Eng *et al*., 2019) and spatial factors were sampled from a Gaussian process with zero mean and Matérn kernel (smoothness parameter *v* = 1.5, lengthscale *ℓ* = 0.5) on a 50 × 50 grid in the unit square followed by a softplus transformation to positive values, normalization to a maximum value of 1 and masking of values outside the cell. Weights were simulated from a standard normal distribution, followed by a softplus transformation to positive values, normalization to a total weight of 1 for every gene and multiplication with independent Bernoulli variables (*p* = 0.7) to induce sparsity. To examine the effect of varying molecule abundance in the data, e.g. caused by differences in detection efficiency or biological differences, we repeated the simulation with different scale factors for the intensity function (*μ_dm_* = 50, 100, 200, 300), resulting in an average of 11, 22, 44 and 66 molecules, respectively, per gene and cell.

Across all simulation scenarios, FISHFactor shows a good recovery of the simulated factors and weights (measured using Pearson correlation between inferred and simulated factors and weights, Fig. 3a) with increasing accuracy for data sets with higher number of molecules. In comparison to NMF and NSF, FISHFactor achieves a better factor correlation in all scenarios. Accuracy of the inferred weights is similar or superior in FISHFactor compared to NMF and NSF, whereby NMF and NSF show a strong sensitivity to the choice of the binning resolution, which needs to be chosen in an optimal manner to reach the accuracy of FISHFactor. Such a choice can be difficult to make on real data, where no ground truth is available, and is not required in FISHFactor. At the same time, FISHFactor provides a more accurate weight reconstruction than NMF and NSF at higher spatial resolutions of factors, while for NMF and NSF accuracy in the weight reconstruction comes at the cost of a lower resolution (Fig. 3a, illustrated for a single cell in Fig. 3c). Notably, NSF fails to converge in more challenging scenarios with small numbers of molecules (fraction of cells with convergence, Fig. 3b) and accuracy estimates for NSF based on the subset of cells with convergence in Fig. 3a might be overly optimistic in these scenarios.

**Figure 3:**
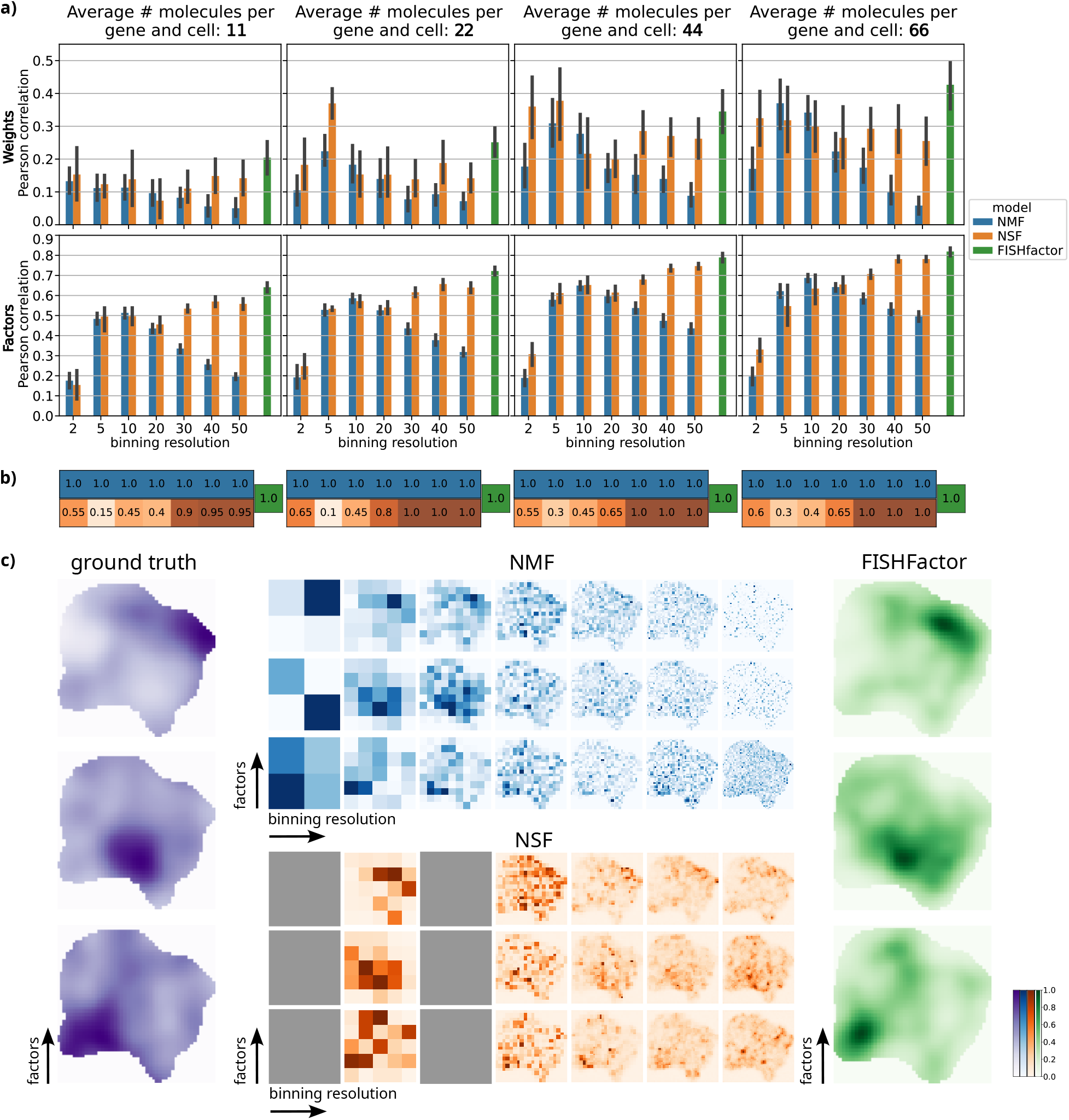
Comparison of FISHFactor to NMF and NSF at different binning resolutions on 20 simulated cells with 4 different intensity scale factors. **a)** Reconstruction accuracy of the simulated weights (first row) and factors (second row). Barplots show the mean Pearson correlation across the 20 cells for FISHFactor and NMF and all cells with convergence for NSF (see b)). Error bars indicate one standard deviation of the mean. **b)** Fractions of included cells (out of 20) for every intensity scale factor and binning resolution. NSF did not converge on all cells. **c)** Exemplary visualization of ground truth and inferred factors in a single cell with an average of 66 molecules per gene. NSF did not converge on the 2 × 2 and 10 × 10 binning resolutions.

### 3.2 Joint modeling of cells improves reconstruction of weights and factors

In a second experiment, we investigated whether for related cells the ability of FISHFactor to share information across cells by jointly modeling their subcellular patterns benefits the reconstruction of weights and factors. For this, we simulated 10 data sets with 20 cells each as described in Subsec. 3.1 using a single shared weight matrix for all 20 cells and an intensity scale factor *μ_dm_* = 100, and applied FISHFactor to this data separately for each cell or jointly modeling multiple cells. We found that the inclusion of multiple cells in the model greatly improves the reconstruction accuracy of the shared weights and, to a smaller extent, the accuracy of the inferred cell-wise factors (Fig. 4).

**Figure 4:**
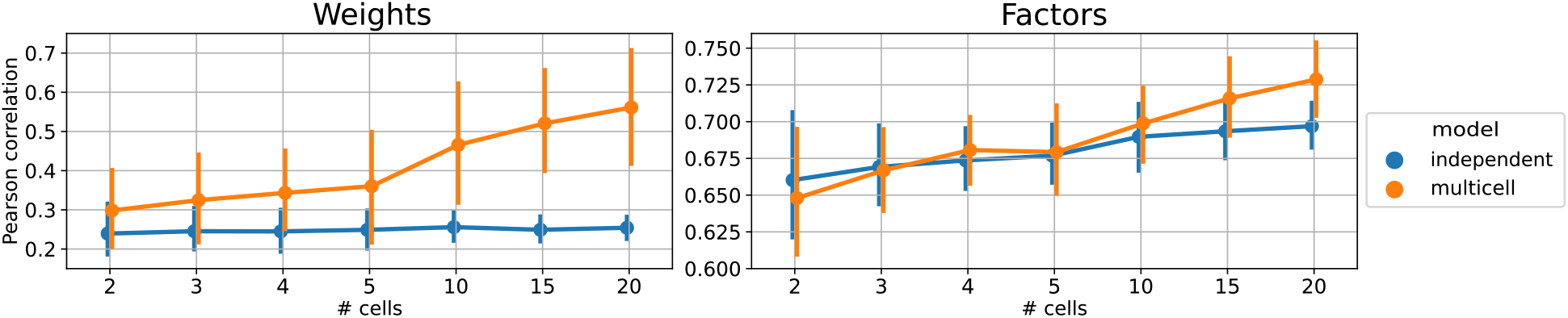
Pearson’s correlation of inferred and simulated weights and factors in 10 simulated data sets of 20 cells with shared weight matrices. In every data set, a given number of cells (x-axis) were modeled using FISHFactor on all cells jointly (multi-cell, orange) or by applying FISHFactor to individual cells and averaging the results (independent, blue). Error bars show one standard deviation of the mean across the 10 data sets.

### 3.3 FISHFactor reveals major gene clusters and subcellular expression patterns in 3T3 cells

Lastly, we applied FISHFactor to a real data set that comprises single-molecule data for 10 000 genes in 225 segmented cultured mouse embryonic fibroblasts (NIH/3T3) from a seqFISH+ experiment (Eng *et al*., 2019). As input for FISHFactor we used all cells and considered genes with a minimum of 30 molecules on average across cells, resulting in a total of 104 genes. Exemplary molecule coordinates for 4 genes in 4 cells are shown in Fig. 5a.

**Figure 5:**
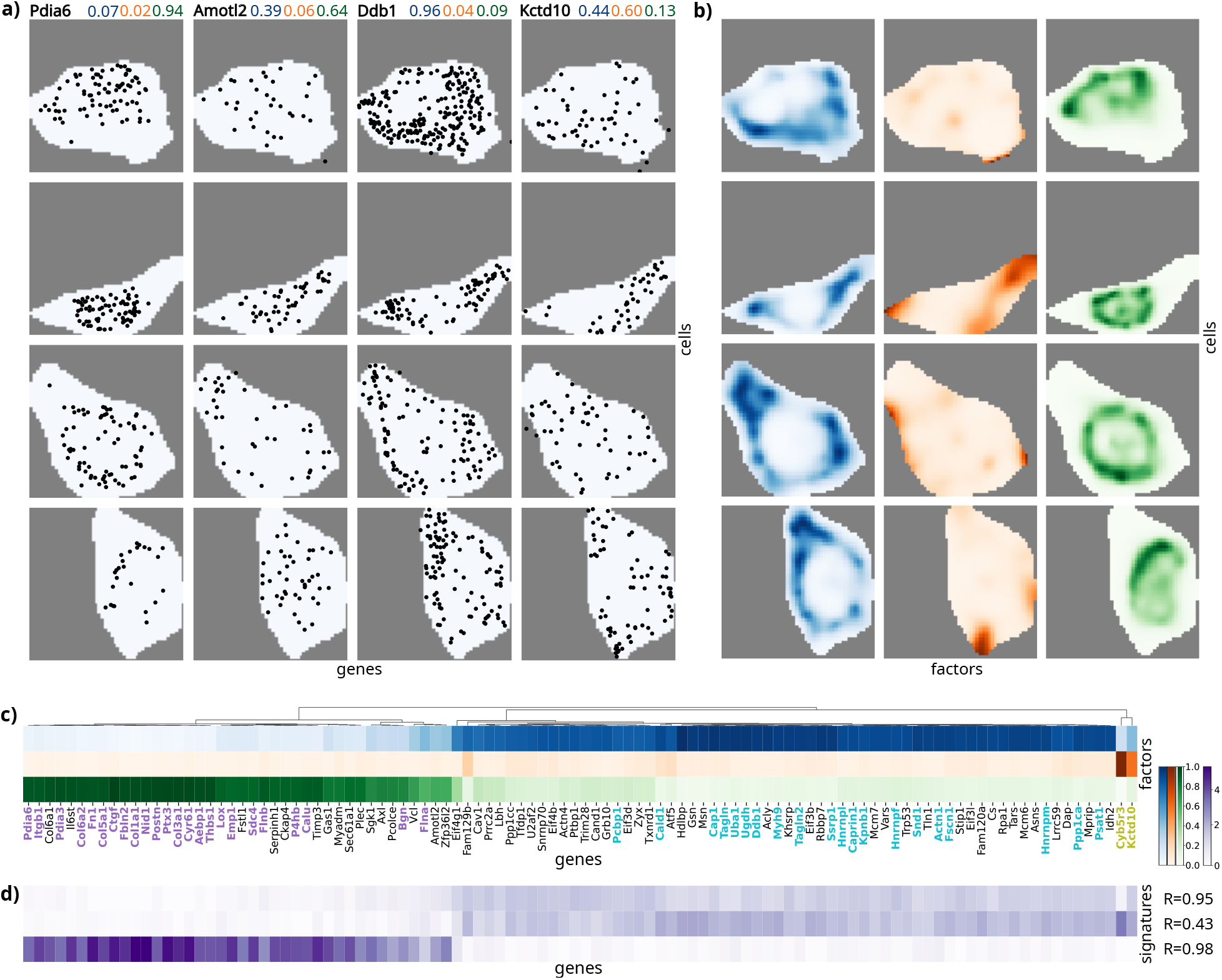
Application of FISHFactor to a data set from cultured mouse embryonic fibroblasts (NIH/3T3) (Eng *et al*., 2019). **a)** Molecule coordinates in 4 cells (rows) for 4 genes (columns) with inferred weights shown in matching colors (blue: factor 0, orange: factor 1, green: factor 2). Pdia6 has high weight on factor 2, Amotl2 on factor 0 and 2, Ddb1 on factor 0 and Kctd10 on factor 1. **b)** Visualization of 3 factors inferred in the same cells. Factor 0 is active around the cell center, factor 1 at the cell border and factor 2 in the cell center. **c)** Hierarchical clustering of inferred gene weights recovers known gene clusters from Eng *et al*. (2019), indicated by label colors (purple: nucleus/perinucleus, cyan: cytoplasm, olive: protrusions). **d)** Gene loadings for signatures identified in Mah *et al*. (2022) correlate strongly with inferred weights in c), the displayed value is the Pearson correlation with the FISHFactor weights.

From these data, FISHFactor identified 3 factors capturing major subcellular expression patterns (Fig. 5b). The factors show distinct subcellular activities, with Factor 0 mainly being active around the cell center, Factor 1 at the cell border and Factor 2 inside the cell center. The inferred weight matrix (Fig. 5c) shows a clear clustering of genes into 3 clusters, where Factor 0 has high weights for genes previously annotated to cytoplasm (Eng *et al*., 2019) (Fig. 5c, cyan), Factor 1 for genes previously annotated to protrusions (Eng *et al*., 2019) (Fig. 5c, olive) and Factor 2 for genes previously annotated to nucleus / perinucleus (Eng *et al*., 2019) (Fig. 5c, purple). We compared this gene clustering with a clustering based on inferred weights of NMF applied to normalized and transformed gene counts per cell, and found that the clustering differs substantially, allowing the conclusion that the spatial information is needed to reconstruct the clusters in Eng et al. (2019) (Fig. S1).

Finally, we asked to what extent the unsupervised approach of FISHFactor is able to recover gene loadings for signatures previously identified on this data based on manual annotation of genes to cellular regions (Mah *et al*., 2022). For this, we compared the factor weights inferred by FISHFactor to the previously identified signature loadings for the same set of genes (Fig. 5d) and found a strong correlation. This indicates that FISHFactor is able to retrieve the same information, but in a completely unsupervised manner. Moreover, the FISHFactor weights appear to be more sparse compared to the signature loadings from Mah et al. (2022), where loadings for signature 0 strongly correlate with loadings for signature 1 (Fig. 5d).

Overall, this application demonstrates that FISHFactor can reveal the major subcellular localization patterns in a data-driven manner without the need for manual labeling or segmentation of areas within the cell and identifies relevant gene clusters based on their subcellular colocalization.

## 4 Discussion

The spatial localization of individual RNA molecules in a cell has long been limited to only a handful of genes at a time. However, recent technological developments have dramatically increased the number of genes that can be profiled, thereby enabling single-molecule resolution of spatial transcriptomics. This opens up the application of computational methods that share information across several genes, such as matrix factorization, which is based on the assumption that spatial expression densities of genes can be linearly decomposed into a small number of initially unknown patterns. While the benefits of such approaches have been demonstrated for spatial transcriptomics data on the cellular level Velten et al. (2022); Townes and Engelhardt (2021), it was unclear how and whether similar ideas could be used to gain insights into the localization patterns of individual molecules at the subcellular level.

Here we addressed this question by developing FISHFactor, a spatial non-negative factor model for singlemolecule resolved spatial transcriptomics data that facilitates the identification of major subcellular expression patterns and co-localization of genes. We demonstrated that the use of a tailored likelihood model for single-molecule data based on a Poisson point process is beneficial compared to naive application of existing factor models that require data aggregation via binning. In addition to sharing information across all genes, FISHFactor furthermore enables sharing information across cells by jointly modeling expression patterns in hundreds of cells, which could improve the reconstruction accuracy of the weights and factors in our simulation studies and provides a direct means to compare expression patterns across cells. A joint modeling of cells can be particularly useful when the number of detected molecules per cell is relatively low and a single cell is not sufficient to reliably identify colocalization patterns of genes. We demonstrated the value of FISHFactor for the unsupervised analysis of single molecule resolved data by an application to a data set of cultured mouse embryonic fibroblasts, where the method identified relevant subcellular expression patterns and gene clusters based on subcellular spatial colocalization. Notably, these clusters cannot be detected on the cellular level, underlining the importance of considering subcellular information.

While memory requirements of FISHFactor are constant in the number of cells, the requirements scale linearly with the total number of molecules per cell and number of factors, which can be a limiting factor for large-scale applications. For example, this limits the number of molecules to approximately 10 000 to 20 000 per cell when using 3 factors on a typical GPU with 10 to 20 GB memory, but future extensions of the model could address this by implementing subsampling strategies on the level of genes or molecules and developing approaches for an automated choice of relevant genes. Importantly, the current implementation of FISHFactor relies on having accurate cell segmentations available to assign molecules to cells. For future research it would therefore be interesting to investigate the benefits of joint segmentation and modeling approaches. Moreover, lifting the restriction of a single shared weight matrix for all cells and instead allowing a priori unknown groups of cells to share group-specific weight matrices would make the model even more flexible for heterogeneous cell populations with different gene colocalizations.

## Data availability

The data used in this article were accessed from Zenodo, at https://doi.org/10.5281/zenodo.2669683; and from figshare, at https://doi.org/10.6084/m9.figshare.15109236.v2.

## Acknowledgments

The authors thank William Townes for his support for using the NSF implementation, and the Pyro and GPyTorch teams and communities for assistance in implementing FISHFactor.

## Supplementary materials

**Figure S1:**
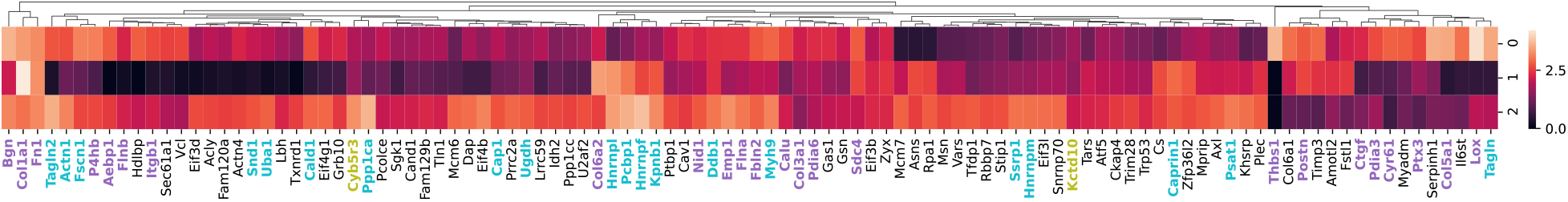
Gene clustering based on total RNA counts per cell, without including spatial information. Gene counts were normalized by dividing by the total counts per cell (considering all 10 000 genes) and transformed as log(10000x + 1) where *x* is the normalized gene count. NMF with 3 components was performed with the normalized and transformed gene counts and genes were clustered using hierarchical clustering on the weights. Label colors denote clusters identified in Eng *et al*. (2019) (purple: nucleus/perinucleus, cyan: cytoplasm, olive: protrusions).

